# Neuronal-spiking-based closed-loop stimulation during cortical ON and OFF states in freely moving mice

**DOI:** 10.1101/2022.02.28.482319

**Authors:** Martin Kahn, Lukas B. Krone, Cristina Blanco-Duque, Mathilde C.C. Guillaumin, Edward O. Mann, Vladyslav V. Vyazovskiy

## Abstract

The slow oscillation (SO) is a central neuronal dynamic during sleep and is generated by alternating periods of high and low neuronal activity (ON and OFF states). Mounting evidence causally links the SO to sleep’s functions, and it has recently become possible to manipulate the SO non-invasively and phase-specifically. These developments represent promising clinical avenues, but they also highlight the importance of improving our understanding of how ON/OFF states affect incoming stimuli and what role they play in neuronal plasticity. Most studies using closed-loop stimulation rely on the electroencephalogram (EEG) and local field potential (LFP) signals, which reflect neuronal ON and OFF states only indirectly. Here we develop an online detection algorithm based on spiking activity recorded from laminar arrays in mouse motor cortex. We find that online detection of ON and OFF states reflects specific phases of spontaneous LFP SO. Our neuronal-spiking-based closed-loop procedure offers a novel opportunity for testing the functional role of SO in sleep-related restorative processes and neural plasticity.

## Introduction

The possibility of non-invasive modulation of sleep oscillations has recently attracted significant attention (Bellesi, Riedner, Garcia-Molina, Cirelli, & Tononi, 2014; Choi, Kwon, & Jun, 2020; Fattinger et al., 2019; Frase et al., 2019; Geiser et al., 2020; Krugliakova et al., 2022; Malkani & Zee, 2020; Marshall, Helgadottir, Molle, & Born, 2006; Ngo, Martinetz, Born, & Molle, 2013; Schneider, Lewis, Koester, Born, & Ngo, 2020). Slow waves are a predominant type of sleep oscillatory activity during NREM sleep, but can also occur during REM sleep and wakefulness (Andrillon, Burns, Mackay, Windt, & Tsuchiya, 2021; Bernardi et al., 2019; Borbely, Tobler, & Hanagasioglu, 1984; Funk, Honjoh, Rodriguez, Cirelli, & Tononi, 2016; Vyazovskiy et al., 2014; Vyazovskiy et al., 2011). Sleep slow waves are homeostatically regulated (Achermann, Dijk, Brunner, & Borbely, 1993; Borbély, 1982; Huber, Deboer, & Tobler, 2000; Krone et al., 2021; Thomas, Guillaumin, McKillop, Achermann, & Vyazovskiy, 2020), and have been implicated in synaptic plasticity, metabolic restoration, glymphatic clearance and other functions (Frank & Heller, 2019; James M. Krueger, Frank, Wisor, & Roy, 2016; Vyazovskiy & Harris, 2013). Traditionally, online detection of slow waves relies solely on their cortical surface- or scalp-recorded electroencephalogram EEG waveforms (Moreira et al., 2021; Ngo et al., 2013; Santostasi et al., 2016), where the specific phase is assumed to correspond to periods of high or low neuronal activity (ON and OFF states) or transitions between population activity and silence (McKillop et al., 2018; Nir et al., 2011). However, in such studies no attempts have been made to directly target the underlying neuronal network activity itself.

The central aim of this study was to develop and validate the methodology for online detection of ON and OFF periods, and to investigate the possibility of neuronal-spiking-based closed-loop stimulation during spontaneous sleep in mice. The potential applications of this method include addressing the following questions:

1. The role of sleep in synaptic plasticity. In vitro evidence and experiments in anesthetized animals suggest that pairing synaptic inputs with population ON and OFF periods leads to plastic changes in neural responses to stimulation (Bartram et al., 2017; Gonzalez-Rueda, Pedrosa, Feord, Clopath, & Paulsen, 2018). This observation is important, as it suggests that a careful choice of the phase of stimulation could make sleep more restorative, but, alternatively, could also lead to sleep disruption and potentially to the development of maladaptive plastic changes within the thalamocortical circuitry. To this end, a better understanding of the role of ON and OFF periods in neural plasticity, as suggested by previous work, is essential.
2. Effects of ON/OFF states during spontaneous sleep on sensory responsiveness and processing of incoming stimuli (Massimini et al., 2005; Nir et al., 2017; Nir, Vyazovskiy, Cirelli, Banks, & Tononi, 2015; Vyazovskiy, Faraguna, Cirelli, & Tononi, 2009). We argued that this is critical, for example, to develop most efficient and least disruptive stimulation protocols, and to establish whether the properties of induced slow waves differ depending on background activity.
3. Correspondence between neuronal activity and LFP waveforms. Finally, given that individual EEG slow waves vary greatly with respect to their origin, shape, amplitude and spatio-temporal dynamics (Bukhtiyarova, Soltani, Chauvette, & Timofeev, 2019; Massimini, Huber, Ferrarelli, Hill, & Tononi, 2004; Murphy et al., 2009; Nir et al., 2011; Riedner, Hulse, Murphy, Ferrarelli, & Tononi, 2011), targeting those directly with conventional closed-loop paradigms likely leads to many instances when stimulation is delivered during a suboptimal or even undesirable phase of the network oscillation. Arguably, this could influence the outcome of modulation. Therefore, obtaining a better understanding of the correspondence between neuronal activity and EEG/LFP waveforms across cortical layers will provide important refinement, both conceptual and methodological, for the approach used to target sleep slow waves.

## Methods

All experiments were carried out in accordance with the UK Animals (Scientific Procedures) Act of 1986. All animals used in this study were C57BL/6 purchased from Harlan Laboratories and kept on a regular (non-inversed) 12h light/dark cycle. Seven male adult C57BL/6 mice (age at baseline recording 125±8 days, body weight: 29.5±0.8 g) were used for all experiments.

### Implants and surgical procedure

All implants were prepared manually before the surgery. For the frontal and occipital EEG recordings, silver wires were wrapped around blunted skull screws and soldered to a 90-Degree connector (Pinnacle Technology Inc. Lawrence). For the electromyogram (EMG), the end of a silver wire was bent into a U-shape and then twisted, to avoid sharp edges. This was done on two separate wires that were soldered to the above-described EEG head stage. The laminar probe (NeuroNexus Technologies; A1 × 16-3mm-100-703-Z16) has a ground and a reference wires, each soldered to male connector pins, which could then be connected during surgery to female connector pins on the ground and reference screw, respectively. The laminar probe was stained with the dye DiI (DiIC18(3), Invitrogen) before surgery to aid the localisation of the electrode tract (Krone et al., 2021).

To induce anaesthesia, the mouse was exposed to a prefilled chamber with 4% isoflurane in medical oxygen and, once the mouse had lost the righting reflex and approached a breathing rate of approximately 80 min^-1^. The animal was then transferred to a heating pad and 2-3% isoflurane administered through a nose mask at an oxygen flow rate of ∼1-1.5 L/min. After the scalp was shaved and cleaned using iodine and ethanol, the animal was transferred to a stereotaxic frame where isoflurane was administered at a concentration of 0.6-1.2 % at a flow rate of ∼1 L/min throughout the surgery. At this point, Metacam^®^ (meloxicam, 5 mg/kg, Boehringer Ingelheim Ltd., Bracknell, UK), Vetergesic^®^ (buprenorphine, 0.1 mg/kg, Sogeval UK Ltd., York, UK) and Dexamethasone (0.2 mg/kg s.c., Boehringer Ingelheim Ltd., Bracknell, UK) were injected subcutaneously and artificial tears were applied. Once the head was fixed, a rectal probe was inserted to maintain core temperature around 37°C. The scalp was opened, and the straightness of the skull was verified by levelling bregma and lambda and the points 1 mm lateral to Bregma. To minimise the loss of implants, the skull’s surface was roughened using the scalpel and etching gel, and the coordinates for implantation were marked as shown in Figure 1a. The holes for the reference (cerebellum), ground (cerebellum or left occipital) and the two EEG screws (frontal and occipital) were drilled first and the screws were then immediately inserted using a screwdriver. Subsequently, the hole for the bipolar concentric stimulation electrode (Plastics One, see Experimental design below for further information) was drilled and the electrode was carefully and slowly inserted. All screws were then fixed with dental cement SuperBond^®^ (Prestige Dental Products Ltd, Bradford, UK) before a craniotomy was made for the laminar electrode. Once the bone was removed, the dura was carefully rolled back with a syringe tip and the laminar probe was immediately inserted until the last of its 16 contacts was below the cortical surface (Figure 1b). The craniotomy was immediately sealed with a silicone gel (KwikSil; World Precision Instruments, Sarasota, FL, USA). The entire head stage was then cemented and the EMG wires inserted into the neck before the skin was sutured, if necessary. Animals were given subcutaneous saline injections following the surgery. Following the surgery animals were carefully monitored at least once a day for seven days and analgesics were administered orally or subcutaneously, if necessary.

**Figure 1:**
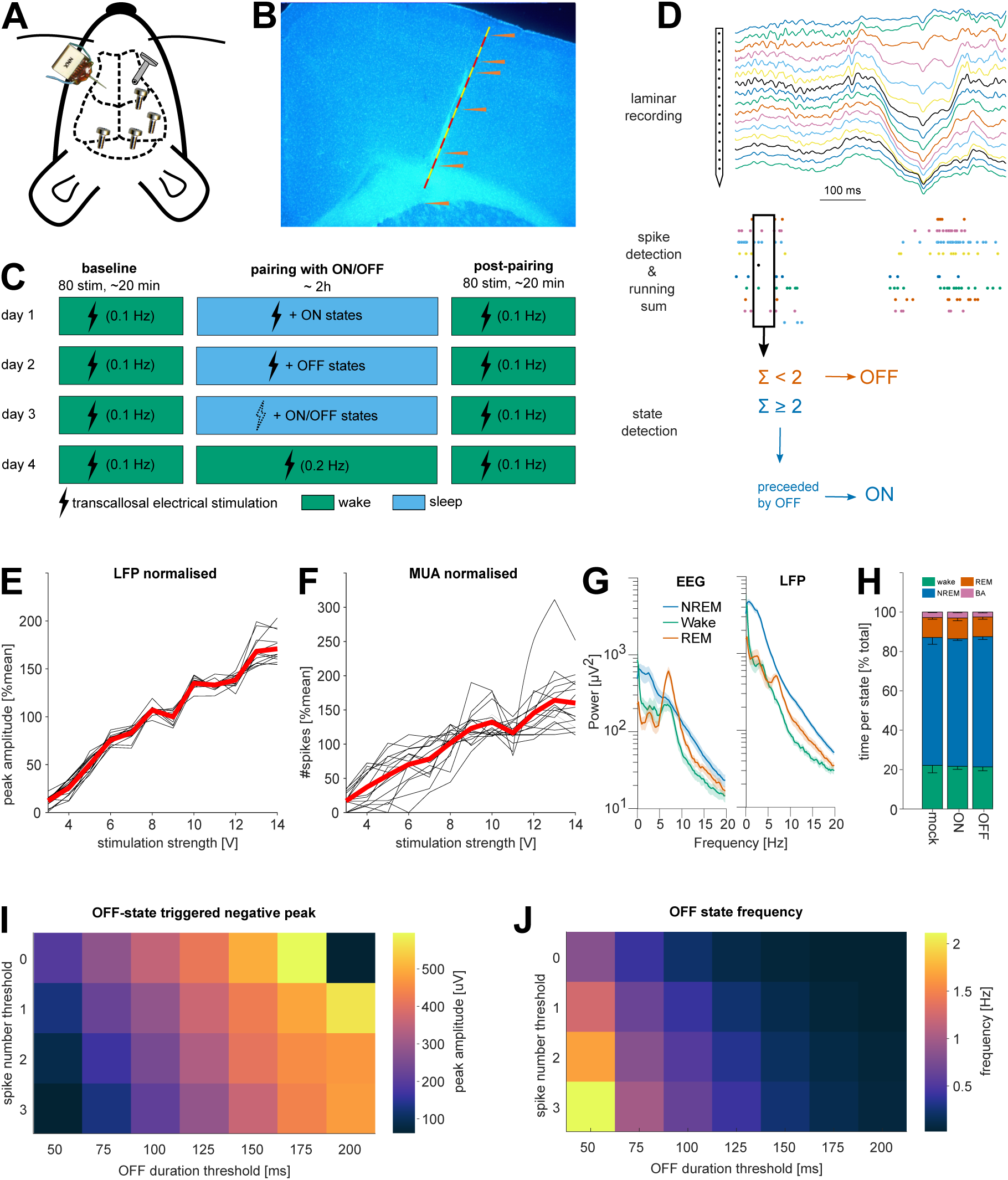
Methodological approach. A. Schematic of cranial electrode placement. 16 channel laminar probe (left) was inserted into M1 contralateral to the stimulation electrode (right), in addition to EEG electrodes in the frontal and the occipital derivation. B. Representative histology image. C. Experimental design. Each mouse was subjected to 4 stimulation paradigms, each on a separate experimental day. Each stimulation paradigm includes a baseline and post-pairing stimulation period with 80 stimulations at 0.1 Hz during wakefulness. D. The method used for ON and OFF periods detection. E. Example of a dose-response curves in the LFP in response to stimulation in one animal. Black lines show the average response across trials of a single channel in the laminar probe. Red line is average across all channels. F. Same as E but for spiking. G. Example of power spectral density in the EEG (left) and LFP (right) during different vigilance states. H. Time spent per vigilance state (+/-SEM) across all mice in the different experimental conditions. I&J. Simulations illustrating the effects of varying parameters of the real-time ON/OFF state detection algorithm. Longer and stricter (i.e. fewer spikes) neuronal silent periods result in a larger peak in the LFP(I), which occur progressively more rarely (J). Note the resulting tradeoff between parameters resulting in a large LFP peak and parameters resulting in frequent detection events.

### Electrophysiological recordings

Animals were moved to recording chambers at least 3 days before the start of any recording. At least 1 day into this habituation phase was allowed before the EEG head stage was connected to a cable bridging the animal and the pre-amplifier and another day before the laminar probe was connected to the pre-amplifier. EEG and EMG signals were routed via an S-box (Tucker Davies Technologies Inc [TDT], Alachua, FL, USA) to a PZ-5 pre-amplifier (TDT), where they were differentially digitised (relative to the cerebellar screw or the contralateral EMG wire, respectively) at 25 kHz. The signal was then sent to a RZ-2 signal processing system (TDT), which interfaced with the Synapse recording software (TDT). The RZ-2 sampled the signal down to 700 Hz (applying an adequate low-pass filter at 45 % of the final sampling frequency) and stored it at 305 Hz. Whenever possible, the signal was stored in this “raw” form in addition to versions with high pass filters more suitable for online monitoring (i.e. 0.5 and 10 Hz for EEG and EMG, respectively).

The signal from the laminar probe was routed directly to the PZ-5 and sampled at 50 kHz. To obtain continuous LFP data (and limit data size) one version of the signal was down sampled to 305 Hz identical to the EEG signal. For stimulation-evoked activity, a snippet of the LFP was stored at 3.5 kHz whenever the recording software triggered a stimulation. Specifically, the snippet started 500 ms before the stimulation and was 1.5 s long. An important consequence of this procedure is that there may be a small (< 1ms) delay between the time when the software sends out the trigger and when the current is applied by the stimulation box.

To record spiking activity, the laminar signal (at 25 kHz) was filtered between 300 and 3000 Hz and a manual threshold was set for each channel individually. The manual threshold was initially set at least 2 standard deviations from the mean. It was then further increased until the average spike waveform (10 s running window) no longer contained noise. Whenever the signal on a given channel crossed the threshold, the timestamp of threshold crossing and a 1.56 ms long snippet of the signal was stored at 12 kHz. This procedure has the advantage that it strongly reduces the considerable data load of recording 16 channels for days at 25 kHz. On the other hand, it irrevocably discards data, especially given than each channel typically recorded spiking activity from more than one individual neuron. In other words, some spikes are too small to trigger the threshold, while some noisy events or spikes produced by one neuron will trigger it and thus create a 1.56 ms long refractory period where spikes produced by other neurons will be lost. Spiking activity was always processed with WaveClus (Chaure, Rey, & Quian Quiroga, 2018). This software was chosen because it was designed explicitly to perform well on single-channel recordings as well as multi-channel recordings. In contrast, most other spike-sorting algorithms are optimised for polytrode recordings (Chung et al., 2017), where a single unit is recorded on >1 channel. Such cases are likely rare in the present recordings, given the relatively large distance between channels (100 μm).

### Experimental Design

Our experimental design included cortical electrical stimulation during both waking and sleep to investigate a) the immediate effects of stimulation on cortical responses, and b) to address the effects of stimulation on synaptic plasticity (Bartram et al., 2017; Vyazovskiy, Cirelli, Pfister-Genskow, Faraguna, & Tononi, 2008; Vyazovskiy, Olcese, Cirelli, & Tononi, 2013) To this end, every mouse was subjected to at least four basic experimental conditions on separate days (Figure 1c). Each condition began with eighty stimulations at 0.1 Hz (the current pulses were 0.1 ms squared monophasic pulses and the chosen output voltage was normally around 6-10 V) approximately at ZT 1 (one hour after lights on). During this “baseline waking” period, mice were kept awake by providing novel objects. Following this baseline stimulation, animals were exposed to the different experimental conditions (described below) for approximately 2.5 h. This will henceforth be referred to as the “pairing” period, because electrical stimulation was typically paired with a specific state (even though no stimulation may occur in some cases). After this pairing period, a post-pairing wake stimulation followed in all conditions. This post-pairing wake stimulation was always identical to the pre-pairing wake stimulation on all days for the same mouse (very subtle differences in baseline stimulation parameters occur in a few mice, but all variance is between mice, never within mice). As shown in Figure 1c, the four basic conditions were (1) sleep-mock: stimulation was targeted alternatingly at ON/OFF states but the stimulation box was turned off. (2) Sleep-ON and (3) sleep-OFF where stimulation was targeted selectively at ON and OFF states, respectively. (4) wake-stim: the same number of stimuli were delivered as during (2) and (3) but the animal was kept awake with novel objects. The inter-stimulus interval was similarly constrained as during (2) and (3) but was adjusted such that the same number of stimuli were delivered in approximately the same amount of time. The number of stimulations delivered during these pairing protocols were determined by the first experimental day in each animal. The animal was allowed to sleep for up to 2.5 hours and the only constraint on stimulation numbers were the minimum interstimulus interval (10 s), the amount of NREM sleep and the number of ON/OFF detections. Thus, interstimulus interval was sufficiently long to prevent induction of plasticity or over-stimulation, but also sufficiently short to obtain a sufficient number of stimulations for subsequent analysis. The total number of stimulations during the pairing period varied slightly between animals but never within animals (i.e. it never varied between conditions). In all subsequent experimental days, the same number of stimulations were delivered (except for mock stimulation days). Therefore, the total duration of the experiment was kept constant at approximately 2.5 hours, but varied slightly (within ∼ 20 min) between conditions. To avoid a systematic effect of repeated stimulation, the order of the conditions was randomised, except that wake-stim (4) was never done as the first condition, because the number of stimuli delivered during the pairing period was constrained most strongly by the ON/OFF detection algorithm. The stimulation strength was chosen based on the dose-response curve (Figure 1e,f, see below). The stimulation strength was then set as the weakest stimulation strength sufficient to elicit a measurable response.

### Online ON and OFF period detection, and closed loop electrical stimulation

Procedures for online data processing and closed-loop stimulation were custom written in the proprietary object-oriented programming environment supplied by TDT and summarised on Figure 1d. First, the incoming spikes were summed across all channels over a predefined time window (usually 50-125 ms). Whenever this running sum went below a predefined threshold (usually 1 or 2 spikes), an OFF state was registered. An ON state was defined as a period of high firing (10-30 Hz) for a prolonged period of time (same duration as OFF state), following an OFF state. To avoid stimulating during waking, a running root mean squared of the EMG signal was used and a manual threshold was set for it. Electrical stimulation was delivered to the animal through a bipolar concentric stimulation electrode attached to a PSI6x stimulus isolation unit. The stimulus isolation unit was coupled to a stimulation box (Grass Instruments), on which the stimulation parameters could be set manually. Once the parameters were set, the stimulation box could be triggered by means of a transistor-transistor logic (TTL) pulse from the RZ-2 system, which was controlled by the recording software. One day before experiments started, an input-output curve was obtained (Figure 1e,f). The 1.5 s long LFP snippets (sampled at 3-6 kHz) surrounding each stimulation were imported into Matlab using the supplier’s (TDT) Matlab software developing kit. Pre-processing of the snippets involved removing line noise and slow drift using a regression-based algorithm (http://chronux.org/ (Mitra & Bokil 2009)). Specifically, we used the Chronux function *locdetrend*, which applies a least squares fit to a running window (800 ms width, 100 ms steps). Regression-based approaches were chosen to avoid introducing filtering artefacts. For analysis of the peak and slope of the evoked response, potential direct current (DC)-offsets were accounted for by subtracting the mean of the 5 ms preceding the stimulation from the entire snippet for each channel and trial separately.

### Histology

After experiments were completed, animals were deeply anaesthetised with an intraperitoneal injection of Pentobarbital (Euthanal). Once the animal reached deep anaesthesia (as verified by loss of righting, pedal and corneal reflexes) microlesions were performed to aid laminar identification of recording sites (Krone et al., 2021). For microlesions, the laminar probe was connected to an impedance testing device (NanoZ, Plexon Inc), which was used to pass current (10 µA for 10 s) through four equally spaced channels of the laminar probe. The bottom channel was always lesioned first, as the first lesion can damage the other channels. Animals were then transcardially perfused with PBS and 4% paraformaldehyde (PFA) and the head of the animal was then stored in 4% PFA (i.e. the implant was not removed at this point, which improved the quality of histology) and moved into acidified PBS after a few days. Brains were embedded in agarose and cut into 50 µM thick coronal sections. The sections were stained with 4′,6-diamidino-2-phenylindole (DAPI) and imaged using a fluorescence microscope. The sections containing the electrode tract were identified using the red Dil fluoresence and were imaged at 1.6, 2.5 and 5x magnification. The recording location in the rostrocaudal and mediolateral dimension was identified using the mouse brain atlas (Paxinos & Franklin, 2001). The cortical layer of each laminar contact was identified in the 5x-magnification images. First, the site(s) of the microlesions were identified in the DAPI, or background fluorescence (green fluorescent protein (GFP)) images. Second, the position of lesions, the Dil staining, and the length of the electrode were used to determine the position of each contact. Layer 1 was identified based on the low density of neurons compared to layer 2/3. Similarly, the beginning of layer 5 was identified based on the lower cell density in layer 5 and the presence of large pyramidal cells characteristic for this layer. While layer 4 is comparatively small in primary motor cortex, it exists and can be identified as a small increase in cell density right above layer 5 (Skoglund, Pascher, & Berthold, 1997; Yamawaki, Borges, Suter, Harris, & Shepherd, 2014). Layer 6 was also identified based on the higher density of cells compared to layer 5.

### Scoring of vigilance states

Data were extracted from the raw data format of the recording software, resampled to 256 Hz and bandpass-filtered using custom Matlab scripts (0.5-100Hz for EEG/LFP and 10-50 Hz for EMG, 3rd order phase conserving type II Chebyshev filter). The signals were then converted to the ASCII format and from there converted into European Data Format (EDF) files. The EDF files were visualised in the software SleepSign (Kissei Comtec Co, Nagano, Japan). To score vigilance states, the LFP, EEG and EMG data were examined in 4-s epochs. If present, timing of electrical stimulation was also visualised. Waking was defined as a low voltage, high frequency EEG with a high variance in the EMG. In contrast, NREM sleep was defined as high amplitude EEG signals containing slow waves (and high delta power) and exhibiting low EMG tone and variance. The EMG commonly displayed clear heartbeat artefacts during all sleep episodes. REM sleep was defined as wake-like activity with sleep-like EMG signal and usually high theta activity in the occipital derivation (resulting EEG and LFP power spectra are shown on Figure 1g). When an animal displayed wake-like activity for less than 4 consecutive epochs (i.e. 16 s) within a NREM bout, this was scored as brief awakening. Episodes of all four types (NREM, REM, waking, brief awakening) were flagged if they contained clear artefacts in any EEG or LFP channel. When sleep scoring was complete, the SleepSign software returned the vigilance states and the power spectra for each 4-s episode; the latter were calculated in 0.25 Hz frequency bins using a Hanning window. Special consideration was given to 4-s epochs containing stimulation events. For stimulations aimed at waking periods, the epoch was only scored as NREM if there was sleep like activity in the 2 s before or after stimulation. Vice versa, if stimulation was aimed at NREM episodes, activity was scored as REM or waking if the activity 2 s before or after the stimulation resembled the respective state. The same “over-sensitive” procedure was applied with regards to artefacts. We found that stimulation did not have major effect on the amount of vigilance states, and >95% of stimulations targeted sleep as intended.

### Statistics

The experimental design of this study posed several statistical challenges. Most notably, each mouse experienced several treatments, and observations were often nested (e.g. multiple channels, per mouse and several mice per condition). To address these challenges, we used linear mixed models (LME) (Harrison et al., 2018). This method has several advantages, most notably it can account for the abovementioned nested nature of experiments and it can readily handle missing data points (e.g. a noisy or unresponsive channel on one day). Each time an LME was used all assumptions of LMEs (independence, homogeneity of variance, normality of error, and linearity) were visually inspected using plots (e.g. QQ plots). To test for significance, we used Matlab and R-studio to fit a model with and without the relevant parameter (e.g. condition) and compared the models using the log-likelihood ratio test. If the result was significant we ran post-hoc Tukey contrast in R-studio.

## Results

### Real-time detection of ON and OFF states during sleep in freely moving mice

We chronically implanted seven mice with frontal and occipital screws to monitor the EEG and with two wires in the neck muscles to measure the EMG. For neuronal activity recording, we implanted a 16-channel laminar probe into the primary motor cortex (M1) (Figure 1a). To detect ON and OFF states online, spikes were summed across all channels of the laminar probe (Figure 1d). OFF states were detected when the running sum of spikes was below a certain threshold (usually below 1 or 2 spikes) for a sufficient amount of time (50-125 ms). An ON state was defined as a period of high firing (10-30 Hz) for a prolonged period of time (same duration as OFF state), following an OFF state. A challenge for this procedure is the trade-off between speed and accuracy and the trade-off between sensitivity and selectivity. Furthermore, the optimal parameters are not uniform across animals, in part due to different numbers of neurons recorded by each laminar probe. Therefore, we used a baseline recording of each mouse to simulate ON/OFF state detection with differing parameters. As expected, increasing the minimum duration of OFF/ON states leads to detection of larger amplitude slow waves in the LFP but also to fewer detections of ON/OFF states (Figure 1i,j), as has been previously reported (McKillop et al., 2018; Vyazovskiy, Olcese, et al., 2009). We surmise that increasing the minimal duration of OFF/ON states leads to an increased chance to detect a state towards its very end.

As expected, we found that OFF state detection was always preceded by a period of neuronal quiescence whereas ON state detections were preceded by increased spiking (Figure 2a). Upon detection of OFF and ON periods, the probability to transition out of the detected state began to increase logarithmically (Figure 2b,c). Importantly, the detection of ON and OFF periods based on neuronal spiking was on average associated with LFP slow waves (Figure 2a), and with expected changes in MUA (Figure 2d). A clear-cut laminar profile of LFP signals associated with detected neuronal ON and OFF periods was apparent (Figure 2e,f), consistent with the notion that LFP slow waves and their underlying neural dynamics originate from the deep cortical layers (Beltramo et al., 2013; Krone et al., 2021; Sanchez-Vives & McCormick, 2000).

**Figure 2:**
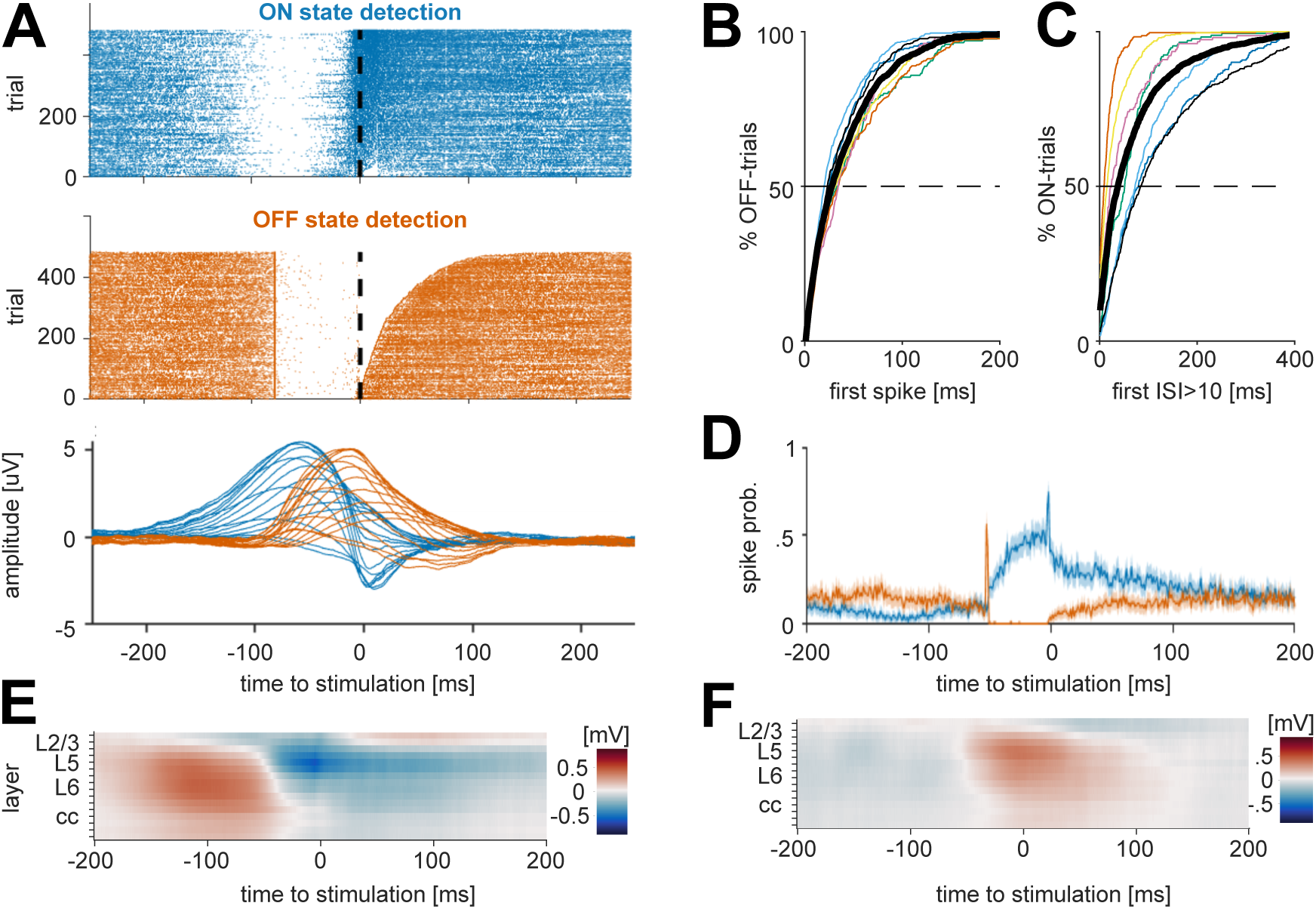
Online detection of ON/OFF states in freely moving mice. A. Top: Example of all ON and OFF state detections in one representative mouse during mock stimulation. Dashed line indicates when stimulation would be delivered. Bottom: Example of average ON/OFF state - evoked LFP signal across all channels in the same animal as shown above. B. Cumulative histogram of the time until the occurrence of first spike following an OFF-state detection. Thin colored lines depict individual mice, thick black line corresponds to the mean across animals. C. Cumulative histogram of the time until the occurrence of first OFF-state following an ON-state detection. Thin colored lines depict individual mice, black line is mean across animals. D. Average ON/OFF state evoked spiking across all channels in one representative mouse. E. Example of average LFP signal across all channels centered on ON-state detections F. Example of average LFP signal across all channels centered on OFF-state detections.

### Neuronal responsiveness differs between ON and OFF periods

For each mouse, we established an input-output curve for electrical stimulation at least one day prior to experiments (Figure 1e,f), and selected the weakest stimulation level that evoked a detectable response in both the MUA and the LFP. We first examined the LFP and MUA response to contralateral stimulation across cortical layers during artefact-free wakefulness epochs (Figure 3a-c). Significant spiking responses (permutation test with 5000 permutations) to electrical stimulation occurred with an average probability of 51 ± 18% (mean ± SD, n=7 mice with 16 channels each) across layers 1, 2/3, 5 and there was a significant effect of layer on response probability (p<0.001, log-likelihood ratio test (dF = 3. χ^2^:37.9)). The spiking response generally involved a period of increased firing, followed by a period where spike rates fell below the spontaneous rates. The increased firing rate began on average 3.66 ± 0.93 ms (mean ± SD; n=72 channels from 7 mice) after stimulation, and started significantly later in layer 1 compared to L5 and L6 (Figure 3b). Notably, in every experiment there was at least one channel that significantly responded within 1 ms of stimulation (mean time to first responsive time bin in any channel across mice: 1.59 ± 0.69 ms (mean ± SD)). This could be due to unaccounted stimulation-induced noise or antidromic activation. The spiking response peaked between 5 and 10 ms, and, on average lasted until 10.7 ± 2.13 ms (mean ± SD) after the stimulus.

**Figure 3:**
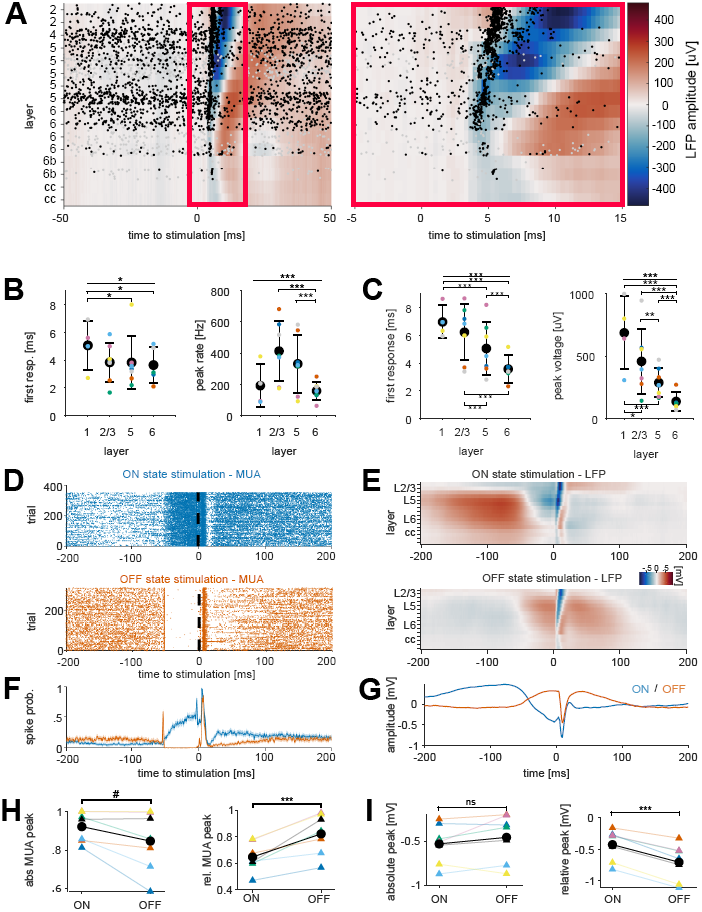
Modulation of evoked responses by stimulation during ON or OFF states. A. Laminar LFP and MUA responses to transcallosal electrical stimulation. Spike occurrences are shown as raster plots. B and C. Characteristics of MUA and LFP responses respectively across cortical layers of M1. *p<0.05, **p<0.01, ***p<0.001. D and E. Representative examples of stimulation-evoked MUA and LFP responses, respectively, during ON (top) and OFF (bottom) states. F and G: Example of average stimulation-evoked MUA and LFPs, respectively, across all channels during ON or OFF states (stimulation delivered at time 0). H and I: Quantification of absolute (left) and normalised (right) stimulation-evoked spike rates and LFP response respectively. #: 0.05 > p < 0.06, ***p<0.001, paired t-test.

The LFP response was closely related to the spiking response but appeared to be slightly delayed. Averaged across all responsive channels, the LFP had a negative peak of 314 µV ± 172 at 8.7 ± 1.85 ms after the stimulation (mean ± SD; n=7 mice with 16 channels each). The LFP response began (defined here as crossing 2 SDs of pre-stimulation baseline) on average 5 ±1.77 ms after the stimulation and lasted for 8.2 ± 1.9 ms (mean ± SD, n= 90 channels from 7 mice), which is consistent with the durations of cortical postsynaptic potentials recorded in single neurons. The anatomical layer had a similar but clearer influence on the LFP compared to the MUA. Similar to the MUA, responsiveness of channels declined with depth, with layer 6 being significantly less likely to respond than all other layers (Figure 3c). Similarly, deep layers had smaller negative peaks, with almost all layer-wise comparisons confirming this pattern. Despite having small peaks, deeper layers tended to respond and peak considerably earlier than superficial layers. Taken together, the layer profile of the evoked response in the MUA and LFP are consistent with a scenario where synaptic inputs reach deep layers first and then reach higher layers through cortico-cortical transmission.

We next examined the evoked responses to ON/OFF stimulation during NREM sleep (Figure 3d-i). The prediction from several previous studies in anaesthetised animals and brain slices is that the magnitude of the response should be significantly modulated by ON/OFF states (Reig et al. 2015; Haider et al. 2007). In line with this, we find that the stimulation-triggered increase in spiking (i.e. relative to pre-stimulation baseline) is larger during OFF-state pairings compared to ON-state pairings (p= 0.0024, paired t-test, n=7). However, when the baseline is not subtracted the opposite trend emerges, with responses during ON states displaying a larger absolute peak (p = 0.0676, paired t-test, n=7, Figure 3h). A similar pattern emerges in the LFP response: when the baseline difference at stimulation onset is accounted for, the response is larger during OFF state stimulation. If it is not, there is no longer any evidence for a difference (Figure 3i).

### Effects of electrical stimulation on sleep architecture and slow wave activity

We next asked whether and how stimulation targeting ON and OFF periods affects sleep. This is relevant because if stimulation during ON and/or OFF states has an immediate effect on sleep (e.g. waking the animal up), then this would be a confound for interpreting the effect of stimulations. However, as shown in Figure 1h, there was no evidence for an effect of stimulation on the relative time spent in NREM sleep, REM sleep or awake (n = 7 mice, 3 separate repeated measures ANOVA, effect of pairing condition on % NREM with sphericity assumed: p=0.518, F(2,12)=0.695; wake: p=0.251, F(2,12)=1.555, or REM (Friedman’s test, p=0.180)).

To test whether stimulation has immediate or delayed effects on the progression of slow-wave activity (SWA) across sleep, we assessed the time course of SWA in the EEG and LFP using only epochs not containing a stimulation event (Figure 4a). As expected, homeostatic decline of SWA resulted in a significant main effect of time on SWA in linear mixed models run separately for the EEG and the average SWA in the LFP (Log-Likelihood Ratio (LLR), χ^2^(9)=112.9 and 147.9 for EEG and LFP, p<10^−16^ for both, n=7 mice). In addition, the parietal EEG displayed a significant effect of condition on SWA (χ^2^(2)=21.62, p=10^−5^), whereas there was no such effect of condition on SWA in the LFP (χ^2^(2)=2.7, p=0.25). There was no evidence for an interaction between condition and time in the EEG/LFP (LLR, χ^2^(18)=16/12, p=0.59/0.8). Post-hoc comparisons in the EEG suggested that OFF state pairings were associated with significantly reduced SWA, compared to ON state pairings and mock pairings (Tukey contrasts, P<10^−5^ for both comparisons in the EEG). This suggests that, independent of when the measurement was taken, the OFF-state pairing condition always displays lower SWA. This contrasts slightly with the visual impression that the first and last time bins are not different and is likely due to insufficient power.

**Figure 4:**
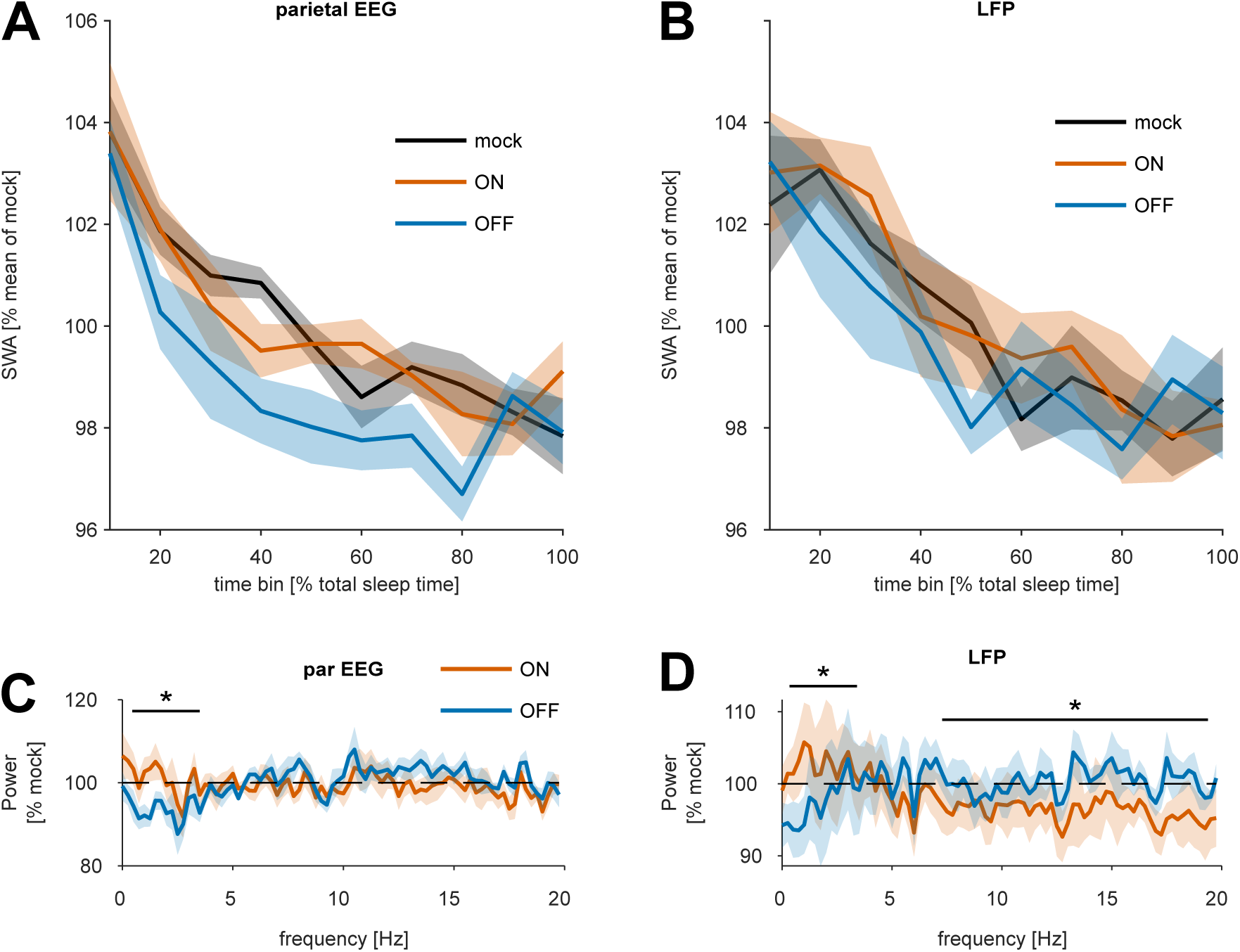
Effects of closed-loop stimulation on sleep EEG. A and B. The time course of EEG slow wave activity (SWA, 0.5-4 Hz) during NREM sleep across the stimulation session in the parietal EEG signal and the LFP recorded from M1 area. Relative values of SWA are plotted in 10-min bins separately for mock stimulation, ON and OFF pairing conditions. Mean values ± SEM, n=7 mice. C and D: The effects of stimulation on EEG and LFP power spectra during NREM sleep. Spectral power values in ON and OFF pairing conditions are expressed as percentage of corresponding frequency bins during the mock stimulation condition for the parietal EEG and LFP signals. Asterisks above denote frequency bands where the effects of ON and OFF pairing differed significantly (p<0.05).

The slow oscillation is not the only network phenomenon during natural NREM sleep, and other events, such as spindles, have been associated with plasticity. To test the effect of stimulation on other frequencies we calculated the difference between the average power spectra (again including only 4-s epochs without stimulation) across conditions (Figure 4c,d). As expected from the previous results, there was a significant interaction between condition and frequency in the LFP (χ^2^(1)=36.9, p=10^−9^, log-likelihood ratio test) and in the frontal EEG (χ^2^(79)=108.17, p=0.016, log-likelihood ratio test). Post-hoc test on individual 0.25-Hz frequency bins suggested that EEG power in the frequencies between 0.25 and 2.5 Hz was lower during OFF-compared to ON-state pairings and not significantly different in other bands (pairwise contrasts without correction for multiple comparisons, bins with p<= 0.01: 0.75-2 Hz, all other significant bands are 0.05>p>0.01, estimated differences ranged from 7 ± 4.11 % (2.5 Hz) to 12 ± 4.11 % (1.25 Hz)). In the LFP (Figure 4d) there was also evidence that ON and OFF pairing had differential effects on power spectra (interaction between condition and frequency: χ^2^(79)=685.8, p=10^−16^, log-likelihood ratio test, Linear mixed effects model (LME) with channels and mouse as nested random effects). Post-hoc tests for individual frequency bins indicated that sleep during ON-state pairings had more power compared to sleep during OFF-state pairings in low frequency bands (0-2 Hz, p<0.001 for all bins except 1.5 Hz with p=0.005), but it had lower power in several higher frequency bands (pairwise contrasts without correction for multiple comparisons, 5.25 – 5.75 Hz, p<0.01; 7 - 8.25 Hz, p<0.05; 9.25 – 9.75 Hz, p<0.05; 10.5 – 20 Hz, p<0.01 for most bins. Together, these data indicate that ON- and OFF-state pairings have a differential effect on the power spectra of NREM episodes that do not contain a stimulation event. Interestingly this is the case for both the LFP and the EEG. Furthermore, this difference is likely driven by the OFF-state pairings, which decrease several frequencies in the SWA range and increase (fewer) frequencies in the spindle range.

### Using closed-loop ON/OFF stimulation to estimate effect sizes of sleep-dependent plasticity

One important application of the approach we describe here is to address the hypothesis that pairing an input to cortex with ON and OFF states has differential effects on synaptic strength. To this end, we recorded LFP and neuronal responses to contralateral electrical stimulation in awake mice exploring objects, and used the magnitude of this response in the LFP and MUA as a proxy for synaptic strength (Fisher et al., 2016; Vyazovskiy et al., 2008; Vyazovskiy, Olcese, et al., 2009). We delivered 80 stimulations (0.1 Hz) before and after each of the following different pairing protocol shown in Figure 1c: stimulation during ON states, stimulation during OFF states, mock stimulation (stimulation turned off), or during waking (novel objects were given to promote wakefulness when necessary).

The effect of the 4 different pairing conditions (referred to as “condition”) on the change in LFP peak amplitude from pre- to post-pairing wakefulness (Figure 5) was assessed with linear mixed-effects models of the form: Δ*V* = *condition* + *V*_*baseline*_ + *condition* * *V*_*baseline*_ + (1|channel: mouse) + (1|mouse). The model supported a significant effect of condition on the change in LFP peak amplitude (Figure 5a). This was true for both the relative change (e.g. V_post_/V_pre_) and the absolute change (e.g. V_post_ - V_pre_) (Figures 5a-b). However, post-hoc tests only yielded significant differences for the relative change, suggesting ON pairings are associated with a stronger decrease in amplitude compared to all other pairings save OFF-state pairings (Tukey-adjusted contrasts for difference in β values ± SE: ON-mock: -7.2 % ± 2.4, p=0.03; ON-wake: -10.39 % ± 2.4, p<0.001; ON-OFF: -4.71 % ± 2.4, p=0.37) (Figure 5a). There was no evidence for a significant interaction between baseline amplitude and condition. While wake pairings had a trend towards increasing the response, this was not significant. We applied the same statistical analysis to the neuronal firing rates (Figure 5c,d). In contrast to the LFP response, the model did not support an effect of stimulation on the relative change in response (Figure 5c). While the model supported an effect of condition on the absolute change in the number of spikes in response to stimulation (Figure 5d), no post-hoc test was significant. Visual inspection of the data suggested that the condition with the biggest effect was pairing of stimulation with wakefulness.

**Figure 5:**
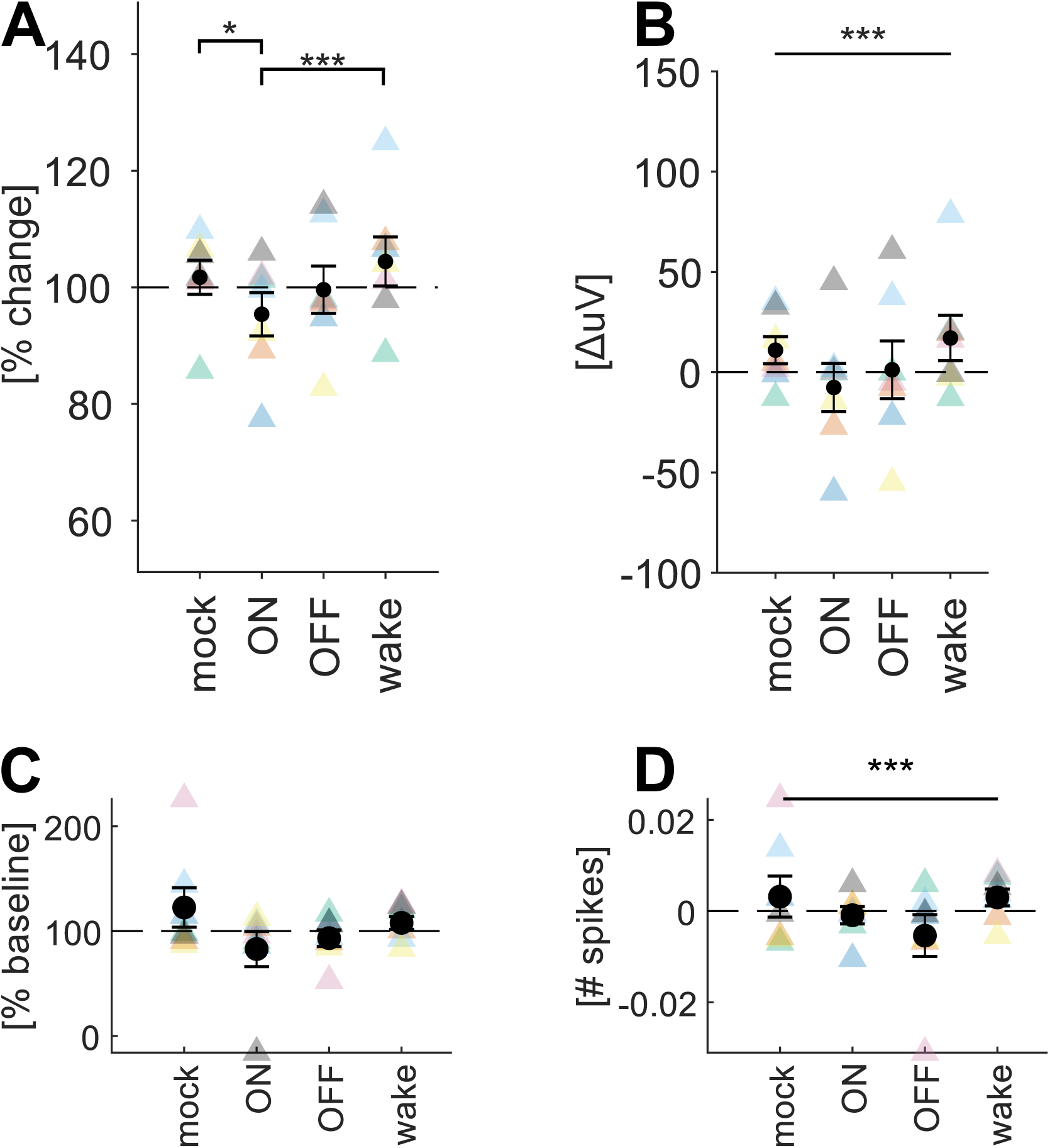
Effects of stimulation targeting ON and OFF periods during sleep on MUA and LFP responses during waking. A. Changes in average LFP amplitude triggered by stimulation during wakefulness before and after pairing stimulation with ON or OFF states. Peak responses were normalised by their baseline amplitude. Black circles show mean +/-SEM and coloured triangles show individual mice. Bars represent pairwise comparisons. *p<0.05, **p<0.01, ***p< 0.001. B. Same as in A but without normalisation to baseline. Bars spanning all conditions indicates significant main effect of condition. ***p< 0.001. C and D. Same as in A&B but for MUA.

In summary, our data suggest that different neuronal states have varying effects on neuronal plasticity. However, using the methodology in the present paper, the differences were subtle (around 5%), and would thus require a substantially bigger sample size to allow more robust conclusions.

## Discussion

Here we developed a method for online detection of cortical ON/OFF states during spontaneous sleep in freely-behaving laboratory mice. While closed-loop stimulation during slow waves is becoming increasingly popular, studies are typically based on the LFP or EEG signals only (Bellesi et al., 2014; Fattinger et al., 2019; Ngo et al., 2013; Schneider et al., 2020), which only indirectly reflect underlying network ON and OFF states (Thomas et al., 2020; Timofeev, 2013). One biologically effective means for closed-loop stimulation relies on setting an (adaptive) negative threshold to detect presumed OFF states and then targeting stimuli into the subsequent UP state by using the average delay between positive and negative peak for each individual (Ngo et al. 2013). This method affects memory (Ngo et al., 2013), changes SWA, and influences the immune system (Besedovsky et al., 2017). However, no studies until now have undertaken a direct online targeting of neuronal network ON and OFF states during sleep.

Our key conclusion is that online targeting of ON and OFF periods based on spiking activity results in a reliable detection of specific phases of LFP slow waves. Our study is consistent with the existing knowledge that spontaneous LFP and EEG slow waves, on average, correspond to a general reduction or a complete cessation of neural spiking, reflecting population OFF periods. It should be noted, however, that sleep has properties of a local process (J. M. Krueger et al., 2008), and, arguably, the neocortex is never entirely in an ON or OFF period (Nir et al., 2011; Siclari & Tononi, 2017; Timofeev, 2013). Therefore, targeting a specific phase of a slow wave in one cortical region will be likely associated with targeting a different – and thus potentially sub-optimal – phase in a different area of the brain. The consequences of such differential manipulations of slow waves in different cortical areas remain to be determined. We further observed that the evoked responses in the LFP began and peaked in deeper layers before the superficial layers. In the MUA, this trend was much less clear and likely present in only a subset of animals. An early response in deeper layers would be consistent with a strong innervation of layer 5 by callosal projections (Petreanu, Huber, Sobczyk, & Svoboda, 2007). However, such a pattern is also conceivable via polysynaptic pathways (i.e. contralateral M1 – region X – M1) and not least via antidromic activation. And yet, even if the laminar probe were placed perfectly in the area most strongly innervated by the stimulated area, or if only antidromic activation occurred, most of the recorded responses (particularly in the LFP) would likely be the consequence of local synaptic connectivity.

One potential application of our new method is to explore the role of sleep and associated patterns of population neuronal activity in synaptic plasticity, to which the current study provides initial, proof-of-principle results. Numerous studies demonstrated that cortical synaptic strength and firing activity are dynamically modulated across the day, or, more precisely, as a function of sleep-wake cycle (Cirelli, 2017; Hengen, Torrado Pacheco, McGregor, Van Hooser, & Turrigiano, 2016; Seibt & Frank, 2019; Watson, Levenstein, Greene, Gelinas, & Buzsaki, 2016). Sleep was linked with strengthening of some synaptic connections (which is thought to mediate consolidation of long-term memories) (Chauvette, Seigneur, & Timofeev, 2012), and weakening or elimination of others (de Vivo et al., 2017), which is thought to allow homeostatic rebalancing of net synaptic strength across the network (Watson et al., 2016). Evidence supporting the profound effects of sleep-wake states on synaptic plasticity includes LFP correlates, such as changes in slope of population synaptic response to stimulation (Chauvette et al., 2012; Vyazovskiy et al., 2008), neuronal activity (Fisher et al., 2016; Watson et al., 2016), phosphorylation status of receptors (Bruning et al., 2019; Diering et al., 2017; Noya et al., 2019; Vyazovskiy et al., 2008), and ultrastructural evidence obtained using electron microscopy (de Vivo et al., 2017).

It is not yet clear whether changes in firing rates between the awake and sleep conditions are causal for synaptic changes, but our previous work indicates that pairing synaptic inputs with ON states would weaken these inputs more strongly than pairing them with OFF states or waking activity (Bartram et al., 2017). To test this hypothesis, here we used an experimental paradigm modified from (Vyazovskiy et al., 2008), to investigate whether pairing of electrical stimulation with ON and OFF periods during spontaneous NREM sleep leads to plastic changes in the motor cortex. Our preliminary data suggest that in the LFP, ON pairings significantly reduced the evoked responses compared to other conditions except OFF pairings. However, OFF pairings did not significantly reduce the peak amplitude. Hence, this data set does not provide strong evidence that OFF pairings reduced the LFP response amplitude whereas ON pairings did so. The lack of significant difference between ON and OFF pairings could have to do with the incomplete separation between ON and OFF states by the algorithm. Wake pairings were only significantly different compared to ON pairings but showed a clear trend to increase the evoked response compared to other conditions. Indeed, the time of peak was significantly delayed by wake pairings, which could support the notion that wake pairing has a significant effect of its own. In the MUA, ON state pairings and wake pairings seemed to increase baseline firing rates. After correcting for changes in baseline firing, there was a significant effect of condition on the change in the mean number of evoked spikes. However, there was no evidence that ON state pairings led to a weakening. Overall, plasticity in the MUA appeared to be very subtle and did not show any conclusive directionality. We surmise that due to the well-known variability between individual neurons (including the possibility that excitatory and inhibitory synapses are modulated in distinct ways (Bridi et al., 2019)), the required sample size to observe an effect would have to be much larger in the MUA than in the LFP, and would possibly require single unit resolution.

Although our stimulation paradigm elicited only minor changes on sleep and SWA, future studies should consider the possibility of biologically significant effects of closed-loop stimulation on other sleep characteristics, beyond merely sleep oscillations, as well as establish if there are any possible long-term effects. If and how state-specific stimulation modulates sleep is of importance because it could have an arousing (Segundo, Naquet, & Buser, 1955) or sleep-promoting (Akert, Koella, & Hess, 1952) effect, or alter sleep intensity (Landsness, Goldstein, Peterson, Tononi, & Benca, 2011). To assess this, power spectral density was assessed in all 4-s epochs that did not include a stimulation. This analysis revealed that OFF-state pairings significantly decreased SWA activity compared to mock and ON-state pairings, in a manner not linearly dependent on time in both the LFP and EEG. Furthermore, frequencies above approximately 11 Hz had increased power in the OFF-compared to ON-state pairings. This strongly indicates a direct effect of OFF-state pairings on sleep oscillations. The shift from lower to higher frequencies seems more consistent with arousal than with local changes in SWA. Our findings do not fully agree with previous studies using closed-loop stimulation. For example, (Ngo et al., 2013) calculated spectra across all 4-s epochs during the pairing period and found an increase in EEG SWA when auditory stimulation was targeted to the UP state and a decrease when the DOWN state was targeted. However, when epochs including stimulation were excluded from the latter analysis, the effect was no longer evident.

In summary, our study provides important new data demonstrating feasibility of *in vivo* targeting of neuronal OFF and ON periods in mice – the network counterparts of EEG or LFP slow waves. This method does not only represent a proof-of-concept that will inform translational studies, but it also establishes a new model for investigating the functional role of the slow oscillation in offline sensory processing and synaptic plasticity.

## Acknowledgements

This work was supported by the Wellcome Trust PhD studentships 203971/Z/16/Z to LBK and 109059/Z/15/Z to CBD. MCK was supported by a Berrow Foundation Lord Florey Scholarship. MCCG was supported by a BBSRC DTP grant (BB/J014427/1) and by a Clarendon Scholarship (provided by the University of Oxford). This work was further supported by a Wellcome Trust Strategic Award 098461/Z/12/Z, John Fell OUP Research Fund Grant 131/032, and Medical Research Council (UK) grant MR/S01134X/1.

## Author contributions

EOM, MCK and VVV designed the study. MCK, LBK, CBD and MCCG conducted the experiments. MCK analysed the data. LBK and MCK performed histology. MCK and VVV wrote the manuscript with the input from all authors.

## References

Achermann, P., Dijk, D. J., Brunner, D. P., & Borbely, A. A. (1993). A model of human sleep homeostasis based on EEG slow-wave activity: quantitative comparison of data and simulations. Brain Res Bull, 31(1-2), 97–113.

Akert, K., Koella, W. P., & Hess, R. Jr., (1952). Sleep produced by electrical stimulation of the thalamus. Am J Physiol, 168(1), 260–267. doi:10.1152/ajplegacy.1951.168.1.260

Andrillon, T., Burns, A., Mackay, T., Windt, J., & Tsuchiya, N. (2021). Predicting lapses of attention with sleep-like slow waves. Nat Commun, 12(1), 3657. doi:10.1038/s41467-021-23890-7

Bartram, J., Kahn, M. C., Tuohy, S., Paulsen, O., Wilson, T., & Mann, E. O. (2017). Cortical Up states induce the selective weakening of subthreshold synaptic inputs. Nat Commun, 8(1), 665. doi:10.1038/s41467-017-00748-5

Bellesi, M., Riedner, B. A., Garcia-Molina, G. N., Cirelli, C., & Tononi, G. (2014). Enhancement of sleep slow waves: underlying mechanisms and practical consequences. Front Syst Neurosci, 8, 208. doi:10.3389/fnsys.2014.00208

Beltramo, R., D’Urso, G., Dal Maschio, M., Farisello, P., Bovetti, S., Clovis, Y., … Fellin, T. (2013). Layer-specific excitatory circuits differentially control recurrent network dynamics in the neocortex. Nat Neurosci. doi:nn.3306 [pii]10.1038/nn.3306

Bernardi, G., Betta, M., Ricciardi, E., Pietrini, P., Tononi, G., & Siclari, F. (2019). Regional Delta Waves In Human Rapid Eye Movement Sleep. J Neurosci, 39(14), 2686–2697. doi:10.1523/JNEUROSCI.2298-18.2019

Besedovsky, L., Ngo, H. V., Dimitrov, S., Gassenmaier, C., Lehmann, R., & Born, J. (2017). Auditory closed-loop stimulation of EEG slow oscillations strengthens sleep and signs of its immune-supportive function. Nat Commun, 8(1), 1984. doi:10.1038/s41467-017-02170-3

Borbély, A. A. (1982). A two process model of sleep regulation. Hum Neurobiol, 1(3), 195–204.

Borbely, A. A., Tobler, I., & Hanagasioglu, M. (1984). Effect of sleep deprivation on sleep and EEG power spectra in the rat. Behav Brain Res, 14(3), 171–182. doi:10.1016/0166-4328(84)90186-4

Bridi, M. C. D., Zong, F. J., Min, X., Luo, N., Tran, T., Qiu, J., … Kirkwood, A. (2019). Daily Oscillation of the Excitation-Inhibition Balance in Visual Cortical Circuits. Neuron. doi:10.1016/j.neuron.2019.11.011

Bruning, F., Noya, S. B., Bange, T., Koutsouli, S., Rudolph, J. D., Tyagarajan, S. K., … Robles, M. S. (2019). Sleep-wake cycles drive daily dynamics of synaptic phosphorylation. Science, 366(6462). doi:10.1126/science.aav3617

Bukhtiyarova, O., Soltani, S., Chauvette, S., & Timofeev, I. (2019). Slow wave detection in sleeping mice: Comparison of traditional and machine learning methods. J Neurosci Methods, 316, 35–45. doi:10.1016/j.jneumeth.2018.08.016

Chaure, F. J., Rey, H. G., & Quian Quiroga, R. (2018). A novel and fully automatic spike-sorting implementation with variable number of features. J Neurophysiol, 120(4), 1859–1871. doi:10.1152/jn.00339.2018

Chauvette, S., Seigneur, J., & Timofeev, I. (2012). Sleep oscillations in the thalamocortical system induce long-term neuronal plasticity. Neuron, 75(6), 1105–1113. doi:S0896-6273(12)00800-8 [pii] 10.1016/j.neuron.2012.08.034

Choi, J., Kwon, M., & Jun, S. C. (2020). A Systematic Review of Closed-Loop Feedback Techniques in Sleep Studies-Related Issues and Future Directions. Sensors (Basel), 20(10). doi:10.3390/s20102770

Chung, J. E., Magland, J. F., Barnett, A. H., Tolosa, V. M., Tooker, A. C., Lee, K. Y., … Greengard, L. F. (2017). A Fully Automated Approach to Spike Sorting. Neuron, 95(6), 1381–1394 e1386. doi:10.1016/j.neuron.2017.08.030

Cirelli, C. (2017). Sleep, synaptic homeostasis and neuronal firing rates. Curr Opin Neurobiol, 44, 72–79. doi:10.1016/j.conb.2017.03.016

de Vivo, L., Bellesi, M., Marshall, W., Bushong, E. A., Ellisman, M. H., Tononi, G., & Cirelli, C. (2017). Ultrastructural evidence for synaptic scaling across the wake/sleep cycle. Science, 355(6324), 507–510. doi:10.1126/science.aah5982

Diering, G. H. S. N. R.,, Roth, R. H., Worley, P. F., Pandey, A., & Huganir, R. L. (2017). Homer1a drives homeostatic scaling-down of excitatory synapses during sleep. Science, 355(6324), 511–515.

Fattinger, S., Heinzle, B. B., Ramantani, G., Abela, L., Schmitt, B., & Huber, R. (2019). Closed-Loop Acoustic Stimulation During Sleep in Children With Epilepsy: A Hypothesis-Driven Novel Approach to Interact With Spike-Wave Activity and Pilot Data Assessing Feasibility. Front Hum Neurosci, 13, 166. doi:10.3389/fnhum.2019.00166

Fisher, S. P., Cui, N., McKillop, L. E., Gemignani, J., Bannerman, D. M., Oliver, P. L., … Vyazovskiy, V. V. (2016). Stereotypic wheel running decreases cortical activity in mice. Nat Commun, 7, 13138. doi:10.1038/ncomms13138

Frank, M. G., & Heller, H. C. (2019). The Function(s) of Sleep. Handb Exp Pharmacol, 253, 3–34. doi:10.1007/164_2018_140

Frase, L., Selhausen, P., Krone, L., Tsodor, S., Jahn, F., Feige, B., Nissen, C. (2019). Differential effects of bifrontal tDCS on arousal and sleep duration in insomnia patients and healthy controls. Brain Stimul, 12(3), 674–683. doi:10.1016/j.brs.2019.01.001

Funk, C. M., Honjoh, S., Rodriguez, A. V., Cirelli, C., & Tononi, G. (2016). Local Slow Waves in Superficial Layers of Primary Cortical Areas during REM Sleep. Curr Biol, 26(3), 396–403.

Geiser, T., Hertenstein, E., Feher, K., Maier, J. G., Schneider, C. L., Zust, M. A., Nissen, C. (2020). Targeting Arousal and Sleep through Noninvasive Brain Stimulation to Improve Mental Health. Neuropsychobiology, 79(4-5), 284–292. doi:10.1159/000507372

Gonzalez-Rueda, A., Pedrosa, V., Feord, R. C., Clopath, C., & Paulsen, O. (2018). Activity-Dependent Downscaling of Subthreshold Synaptic Inputs during Slow-Wave-Sleep-like Activity In Vivo. Neuron, 97(6), 1244–1252 e1245. doi:10.1016/j.neuron.2018.01.047

Harrison, X. A., Donaldson, L., Correa-Cano, M. E., Evans, J., Fisher, D. N., Goodwin, C. E. D., … Inger, R. (2018). A brief introduction to mixed effects modelling and multi-model inference in ecology. PeerJ, 6, e4794. doi:10.7717/peerj.4794

Hengen, K. B., Torrado Pacheco, A., McGregor, J. N., Van Hooser, S. D., & Turrigiano, G. G. (2016). Neuronal Firing Rate Homeostasis Is Inhibited by Sleep and Promoted by Wake. Cell, 165(1), 180–191. doi:S0092-8674(16)30060-5 [pii] 10.1016/j.cell.2016.01.046

Huber, R., Deboer, T., & Tobler, I. (2000). Effects of sleep deprivation on sleep and sleep EEG in three mouse strains: empirical data and simulations. Brain Res, 857(1-2), 8–19.

Krone, L. B., Yamagata, T., Blanco-Duque, C., Guillaumin, M. C. C., Kahn, M. C., van der Vinne, V., … Vyazovskiy, V. V. (2021). A role for the cortex in sleep-wake regulation. Nat Neurosci. doi:10.1038/s41593-021-00894-6

Krueger, J. M., Frank, M. G., Wisor, J. P., & Roy, S. (2016). Sleep function: Toward elucidating an enigma. Sleep Medicine Reviews, 28, 42–50. doi:10.1016/j.smrv.2015.08.005

Krueger, J. M., Rector, D. M., Roy, S., Van Dongen, H. P., Belenky, G., & Panksepp, J. (2008). Sleep as a fundamental property of neuronal assemblies. Nat Rev Neurosci, 9(12), 910–919. doi:nrn2521 [pii] 10.1038/nrn2521

Krugliakova, E., Skorucak, J., Sousouri, G., Leach, S., Snipes, S., Ferster, M. L., … Huber, R. (2022). Boosting Recovery During Sleep by Means of Auditory Stimulation. Frontiers in Neuroscience, 16. doi:10.3389/fnins.2022.755958

Landsness, E. C., Goldstein, M. R., Peterson, M. J., Tononi, G., & Benca, R. M. (2011). Antidepressant effects of selective slow wave sleep deprivation in major depression: a high-density EEG investigation. J Psychiatr Res, 45(8), 1019–1026. doi:S0022-3956(11)00034-3 [pii] 10.1016/j.jpsychires.2011.02.003

Malkani, R. G., & Zee, P. C. (2020). Brain Stimulation for Improving Sleep and Memory. Sleep Med Clin, 15(1), 101–115. doi:10.1016/j.jsmc.2019.11.002

Marshall, L., Helgadottir, H., Molle, M., & Born, J. (2006). Boosting slow oscillations during sleep potentiates memory. Nature, 444(7119), 610–613. doi:nature05278 [pii] 10.1038/nature05278

Massimini, M., Ferrarelli, F., Huber, R., Esser, S. K., Singh, H., & Tononi, G. (2005). Breakdown of cortical effective connectivity during sleep. Science, 309(5744), 2228–2232.

Massimini, M., Huber, R., Ferrarelli, F., Hill, S., & Tononi, G. (2004). The sleep slow oscillation as a traveling wave. J Neurosci, 24(31), 6862–6870.

McKillop, L. E., Fisher, S. P., Cui, N., Peirson, S. N., Foster, R. G., Wafford, K. A., & Vyazovskiy, V. V. (2018). Effects of Aging on Cortical Neural Dynamics and Local Sleep Homeostasis in Mice. J Neurosci, 38(16), 3911–3928. doi:10.1523/JNEUROSCI.2513-17.2018

Moreira, C. G., Baumann, C. R., Scandella, M., Nemirovsky, S. I., Leach, S., Huber, R., & Noain, D. (2021). Closed-loop auditory stimulation method to modulate sleep slow waves and motor learning performance in rats. Elife, 10. doi:10.7554/eLife.68043

Murphy, M., Riedner, B. A., Huber, R., Massimini, M., Ferrarelli, F., & Tononi, G. (2009). Source modeling sleep slow waves. Proc Natl Acad Sci U S A, 106(5), 1608–1613. doi:0807933106 [pii] 10.1073/pnas.0807933106

Ngo, H. V., Martinetz, T., Born, J., & Molle, M. (2013). Auditory Closed-Loop Stimulation of the Sleep Slow Oscillation Enhances Memory. Neuron. doi:S0896-6273(13)00230-4 [pii] 10.1016/j.neuron.2013.03.006

Nir, Y., Andrillon, T., Marmelshtein, A., Suthana, N., Cirelli, C., Tononi, G., & Fried, I. (2017). Selective neuronal lapses precede human cognitive lapses following sleep deprivation. Nat Med, 23(12), 1474–1480.

Nir, Y., Staba, R. J., Andrillon, T., Vyazovskiy, V. V., Cirelli, C., Fried, I., & Tononi, G. (2011). Regional slow waves and spindles in human sleep. Neuron, 70(1), 153–169. doi:10.1016/j.neuron.2011.02.043

Nir, Y., Vyazovskiy, V. V., Cirelli, C., Banks, M. I., & Tononi, G. (2015). Auditory responses and stimulus-specific adaptation in rat auditory cortex are preserved across NREM and REM sleep. Cereb Cortex, 25(5), 1362–1378. doi:10.1093/cercor/bht328

Noya, S. B., Colameo, D., Bruning, F., Spinnler, A., Mircsof, D., Opitz, L., … Brown, S. A. (2019). The forebrain synaptic transcriptome is organized by clocks but its proteome is driven by sleep. Science, 366(6462). doi:10.1126/science.aav2642

Paxinos, G., & Franklin, K. B. J. (2001). The mouse brain in stereotaxic coordinates (2nd ed.). San Diego: Academic Press.

Petreanu, L., Huber, D., Sobczyk, A., & Svoboda, K. (2007). Channelrhodopsin-2-assisted circuit mapping of long-range callosal projections. Nat Neurosci, 10(5), 663–668. doi:10.1038/nn1891

Riedner, B. A., Hulse, B. K., Murphy, M. J., Ferrarelli, F., & Tononi, G. (2011). Temporal dynamics of cortical sources underlying spontaneous and peripherally evoked slow waves. Prog Brain Res, 193, 201–218. doi:B978-0-444-53839-0.00013-2 [pii] 10.1016/B978-0-444-53839-0.00013-2

Sanchez-Vives, M. V., & McCormick, D. A. (2000). Cellular and network mechanisms of rhythmic recurrent activity in neocortex. Nat Neurosci, 3(10), 1027–1034.

Santostasi, G., Malkani, R., Riedner, B., Bellesi, M., Tononi, G., Paller, K. A., & Zee, P. C. (2016). Phase-locked loop for precisely timed acoustic stimulation during sleep. J Neurosci Methods, 259, 101–114. doi:10.1016/j.jneumeth.2015.11.007

Schneider, J., Lewis, P. A., Koester, D., Born, J., & Ngo, H. V. (2020). Susceptibility to auditory closed-loop stimulation of sleep slow oscillations changes with age. Sleep, 43(12). doi:10.1093/sleep/zsaa111

Segundo, J. P., Naquet, R., & Buser, P. (1955). Effects of cortical stimulation on electro-cortical activity in monkeys. J Neurophysiol, 18(3), 236–245. doi:10.1152/jn.1955.18.3.236

Seibt, J., & Frank, M. G. (2019). Primed to Sleep: The Dynamics of Synaptic Plasticity Across Brain States. Front Syst Neurosci, 13, 2. doi:10.3389/fnsys.2019.00002

Siclari, F., & Tononi, G. (2017). Local aspects of sleep and wakefulness. Curr Opin Neurobiol, 44, 222–227.

Skoglund, T. S., Pascher, R., & Berthold, C. H. (1997). The existence of a layer IV in the rat motor cortex. Cereb Cortex, 7(2), 178–180. doi:10.1093/cercor/7.2.178

Thomas, C. W., Guillaumin, M. C., McKillop, L. E., Achermann, P., & Vyazovskiy, V. V. (2020). Global sleep homeostasis reflects temporally and spatially integrated local cortical neuronal activity. Elife, 9. doi:10.7554/eLife.54148

Timofeev, I. (2013). Local origin of slow EEG waves during sleep. Zh Vyssh Nerv Deiat Im I P Pavlova, 63(1), 105–112.

Vyazovskiy, V. V., Cirelli, C., Pfister-Genskow, M., Faraguna, U., & Tononi, G. (2008). Molecular and electrophysiological evidence for net synaptic potentiation in wake and depression in sleep. Nat Neurosci, 11(2), 200–208. doi:10.1038/nn2035

Vyazovskiy, V. V., Cui, N., Rodriguez, A. V., Funk, C., Cirelli, C., & Tononi, G. (2014). The dynamics of cortical neuronal activity in the first minutes after spontaneous awakening in rats and mice. Sleep, 37(8), 1337–1347. doi:10.5665/sleep.3926

Vyazovskiy, V. V., Faraguna, U., Cirelli, C., & Tononi, G. (2009). Triggering slow waves during NREM sleep in the rat by intracortical electrical stimulation: effects of sleep/wake history and background activity. J Neurophysiol, 101(4), 1921–1931. doi:10.1152/jn.91157.2008

Vyazovskiy, V. V., & Harris, K. D. (2013). Sleep and the single neuron: the role of global slow oscillations in individual cell rest. Nat Rev Neurosci, 14(6), 443–451. doi:10.1038/nrn3494

Vyazovskiy, V. V., Olcese, U., Cirelli, C., & Tononi, G. (2013). Prolonged wakefulness alters neuronal responsiveness to local electrical stimulation of the neocortex in awake rats. J Sleep Res, 22(3), 239–250. doi:10.1111/jsr.12009

Vyazovskiy, V. V., Olcese, U., Hanlon, E. C., Nir, Y., Cirelli, C., & Tononi, G. (2011). Local sleep in awake rats. Nature, 472(7344), 443–447. doi:10.1038/nature10009

Vyazovskiy, V. V., Olcese, U., Lazimy, Y. M., Faraguna, U., Esser, S. K., Williams, J. C., … Tononi, G. (2009). Cortical firing and sleep homeostasis. Neuron, 63(6), 865–878. doi:10.1016/j.neuron.2009.08.024

Watson, B. O., Levenstein, D., Greene, J. P., Gelinas, J. N., & Buzsaki, G. (2016). Network Homeostasis and State Dynamics of Neocortical Sleep. Neuron, 90(4), 839–852. doi:S0896-6273(16)30056-3 [pii] 10.1016/j.neuron.2016.03.036

Yamawaki, N., Borges, K., Suter, B. A., Harris, K. D., & Shepherd, G. M. (2014). A genuine layer 4 in motor cortex with prototypical synaptic circuit connectivity. Elife, 3, e05422. doi:10.7554/eLife.05422

